# Chained structure of dimeric F_1_-like ATPase in *Mycoplasma mobile* gliding machinery

**DOI:** 10.1101/2021.04.06.438750

**Authors:** Takuma Toyonaga, Takayuki Kato, Akihiro Kawamoto, Noriyuki Kodera, Tasuku Hamaguchi, Yuhei O Tahara, Toshio Ando, Keiichi Namba, Makoto Miyata

**Author notes:** Address correspondence to Makoto Miyata. Tasuku Hamagichi: Biostructural Mechanism Laboratory, RIKEN SPring-8 Center, 1-1-1 Kouto, Sayo, Hyogo, 679-5148, Japan.

## Abstract

*Mycoplasma mobile*, a fish pathogen, exhibits gliding motility using ATP hydrolysis on solid surfaces, including animal cells. The gliding machinery can be divided into surface and internal structures. The internal structure of the motor is composed of 28 so-called ‘chains’ that are each comprised of 17 repeating protein units called ‘particles’. These proteins include homologs of the catalytic α- and β-subunits of F_1_-ATPase. In this study, we isolated the particles and determined their structures using negative-staining electron microscopy and high-speed atomic force microscopy. The isolated particles were composed of five proteins, MMOBs 1660 (α-subunit homolog), 1670 (β-subunit homolog), 1630, 1620, and 4530, and showed ATP hydrolyzing activity. The 2D structure, with dimensions of 35 and 26 nm, showed a hexameric ring dimer about 12 nm in diameter, resembling F_1_-ATPase catalytic (αβ)_3_. We isolated the F_1_-like ATPase unit, which is composed of MMOBs 1660, 1670, and 1630. Furthermore, we isolated the chain and analyzed the 3D structure, showing that dimers of mushroom-like structures resembling F_1_-ATPase were connected and aligned along the dimer axis at 31 nm intervals. An atomic model of F_1_-ATPase catalytic (αβ)_3_ from *Bacillus* PS3 was successfully fitted to each hexameric ring of the mushroom-like structure. These results suggest that the motor for *M. mobile* gliding shares an evolutionary origin with F_1_-ATPase. Based on the obtained structure, we propose possible force transmission processes in the gliding mechanism.

**IMPORTANCE:** F_1_F_o_- ATPase, a rotary ATPase, is widespread in the membranes of mitochondria, chloroplasts, and bacteria, and converts ATP energy with a proton motive force across the membrane by its physical rotation. Homologous protein complexes play roles in ion and protein transport. *Mycoplasma mobile*, a pathogenic bacterium, was recently suggested to have a special motility system evolutionarily derived from F_1_-ATPase. The present study isolated the protein complex from *Mycoplasma* cells and supported this conclusion by clarifying the detailed structures containing common and novel features as F_1_-ATPase relatives.

## INTRODUCTION

Mycoplasmas are parasitic bacteria characterized by small cell size, a short genome and lack of a peptidoglycan layer (1–3). Many *Mycoplasma* species exhibit a unique gliding motility, which is necessary for their infection (4–6). *Mycoplasma mobile* glides on solid surfaces at 2.0 to 4.5 µm/s in the direction of a protrusion on one side of the cell (Fig. S1) (5). The gliding machinery is localized to the cell protrusion and can be divided into surface and internal structures (Fig. 1A, upper). The surface structure has approximately 450 repeats of a complex of three large proteins, Gli123, Gli521, and Gli349, inserted into the cell membrane (Fig. 1A, lower) (7–11). Fifty-nm-long leg structures corresponding to Gli349 molecules can be seen jutting out from cell protrusion by electron microscopy (EM) (12). The tip of Gli349 is characterized by a “foot” with an oval structure that can bind to sialylated oligosaccharides (SOs) (13–20). Gli521 and Gli123 serve as the “crank” that transfers force to Gli349 and the “mount” that localizes the other two surface proteins to the gliding machinery, respectively. A working model of the gliding mechanism has been proposed in which the cells are propelled by Gli349 molecules that repeatedly catch, pull, and release SOs on solid surfaces (5, 21–23).

**FIG 1.**
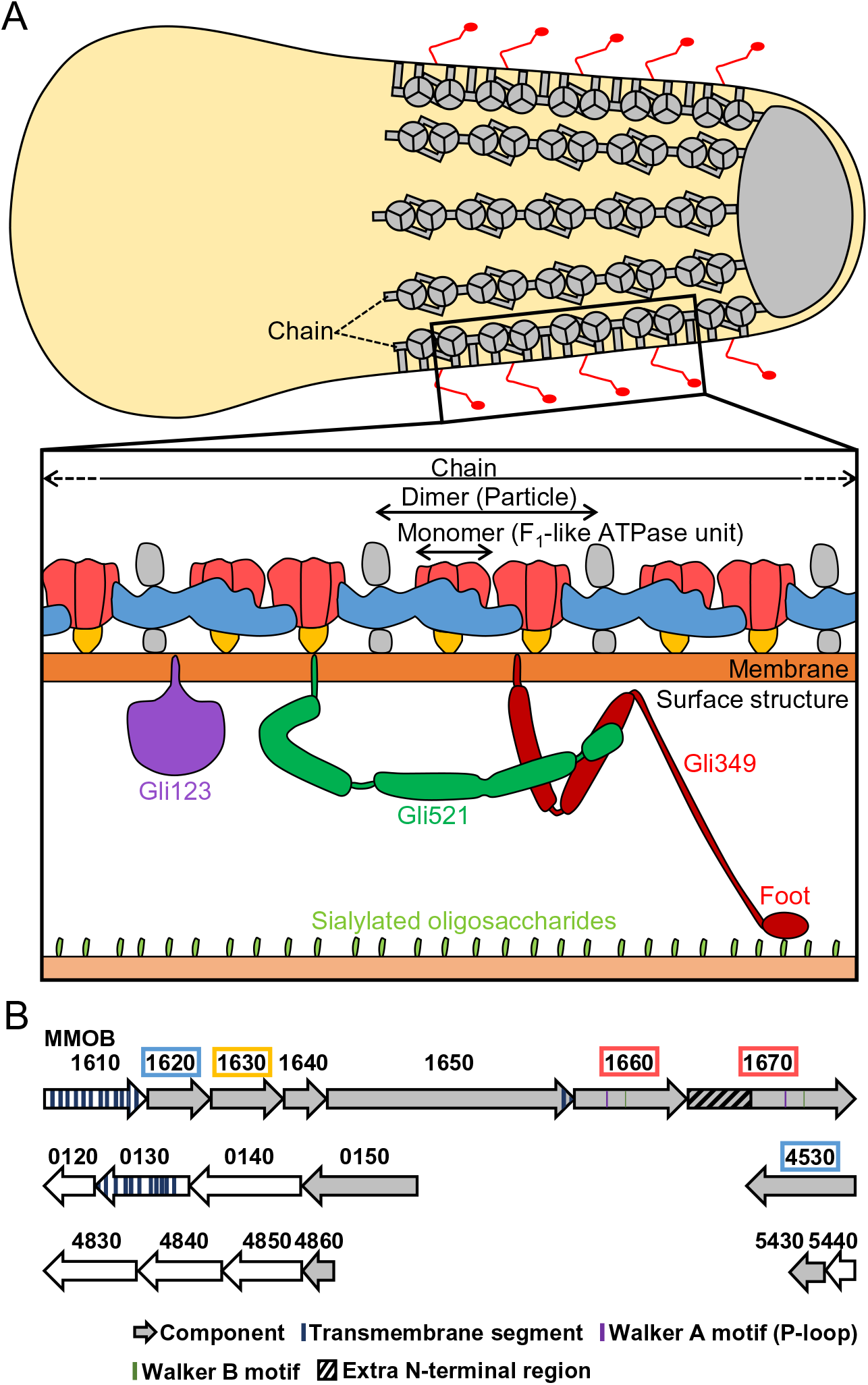
Gliding machinery of *Mycoplasma mobile*. (A) Schematic illustration of gliding machinery based on the present study. In the whole cell shown in upper illustration, the internal structure and legs are colored gray and red, respectively. The actual cell has about 28 chains, each consisting of 17 particles, although a more limited number is illustrated here. In this paper we refer to particle and F_1_-like ATPase unit as the “Dimer” and the “Monomer”, respectively. A single unit of the surface structure and a chain of the internal structure are magnified in the lower illustration. (B) ORFs for the internal structure. The components of the internal structure are colored gray. Type 2 ATPase operon is at the top. Dimer components revealed in the present study are marked by colored boxes, corresponding to the colored components of the lower illustration in panel A.

The internal structure consists of a lumpy structure, called “bell” at the tip of the cell protrusion and 28 “chains” lining the inner membrane surface (Fig. 1A) (23–26). Each chain is characterized by 17 repeating particle structures, resulting in a total of 476 particles in one cell. The chains tend to form sheets when they are isolated from cells, suggesting lateral interaction with the adjacent chains (23, 24). The internal structure consists of least ten proteins (Fig. 1B). Six of these proteins, MMOBs 1620, 1630, 1640, 1650, 1660 and 1670 were coded tandemly in a locus. Interestingly, MMOBs 1660 and 1670, which have Walker A and B motifs that are involved in ATP binding and hydrolysis (27), show high amino acid sequence identity with the catalytic subunits α and β of F_1_-ATPase, respectively. MMOB1670 has an extra N-terminal region (amino acids 1–299), which is not present in the β subunit. The chains most likely contain the motor for gliding, because the gliding motility is coupled to ATP hydrolysis (23, 28, 29).

F_1_F_o_-ATPases, found in most organisms, are rotary motors that perform biological energy conversion (30, 31). Their role is to both synthesize ATP using a proton motive force and, conversely, to hydrolyze ATP to drive protons to maintain the membrane potential. Their structure is composed of a soluble catalytic F_1_-domain for ATP catalysis and a membrane-embedded F_o_-domain for the proton pathway. In the F_1_-domain, the catalytic subunits α and β alternate to form a hexameric ring (αβ)_3_ that rotates the central stalk penetrating the ring using ATP hydrolysis. Phylogenetic studies have shown that *Mycoplasma* have three F_1_-like ATPase clusters, which are referred to as Type 1–3 ATPases (26, 32). Type 1, found in all mycoplasmas, is a typical operon encoding F_1_F_o_-ATPase and is likely to function as a proton pump to maintain membrane potential. Type 3 is found in mycoplasmas that have an MIB-MIP system to cleave host immunoglobulins (33). Type 2 is only found in four *Mycoplasma* species, including *M. mobile*. Interestingly, the Type 2 ATPase of *M. mobile*, which encodes MMOB1620–70, is involved in the internal structure of the gliding machinery.

Recently, the chains of the internal structure were shown to have structural changes linked to ATP hydrolysis. However, it is still unclear how the chain generates and transmits the force to the outside, because its detailed structure has not been clarified. In this study, we isolated the chains and elucidated their structures. The structure had a common architecture with F_1_-ATPase, suggesting that the chain shares a common evolutionary origin with F_1_-ATPase. Based on our findings, we suggest two possible force transmission models for the gliding machinery.

## Results

### Isolation and biochemical analyses of stable unit complex

In this study, we isolated and analyzed three fractions including “Dimer”, “Monomer”, and “Chain” structures which are schematically shown in Fig. 1A (Fig. S2, S3).

First, to isolate a stable unit complex from the chain, we lysed cultured *M. mobile* cells with 1% Triton X-100 and recovered the insoluble fraction by centrifugation (24). This fraction named as “Pellet-1” was used for further preparations. We suspended Pellet-1 in suspension buffer, which contains 137 mM NaCl, and solubilized the putative unit complex for 8 h. Here, we used 137 mM NaCl rather than 400 mM to reduce the contamination of other proteins. The soluble fraction was then subjected to Superdex 200 gel filtration chromatography. The obtained peak fraction in the void contained MMOBs 1620, 1630, 1660, and 1670, which are coded on the mycoplasma Type 2 ATPase operon, and MMOB4530 annotated as phosphoglycerate kinase (PGK) (Fig. 2A). These proteins are known to be components of the internal structure (Fig. 1B) (23–26).

**FIG 2.**
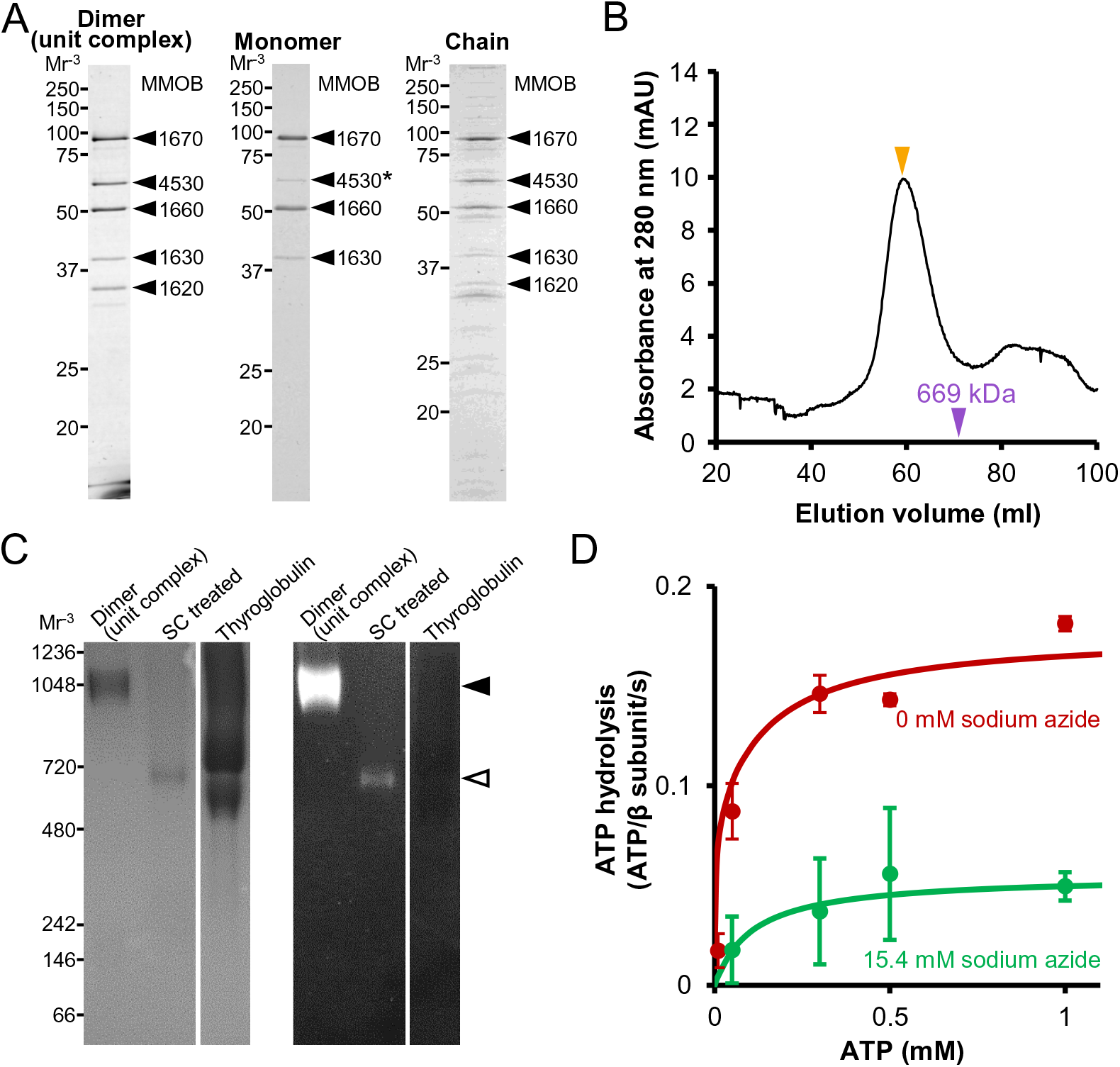
Protein profile and characterization of three fractions. (A) Protein profile of each fraction. Dimer, Monomer, and Chain fractions were subjected to SDS-12.5% PAGE gel and stained with CBB. The bands marked by black triangles were identified by peptide mass fingerprinting (PMF). The intensity of band marked by an asterisk is reduced in Monomer fraction. Molecular masses are shown on the left. (B) Gel filtration assay of unit complex fraction using Sephacryl S-400 HR column. The peak positions of unit complex (Dimer) and thyroglobulin (669 kDa) are marked by orange and purple triangles. (C) BN-PAGE (left) and In-gel ATPase activity assay (right). Dimer, Dimer treated with 1.5% sodium cholate (SC treated), and thyroglobulin, which has no ATPase activity, were subjected to 3 to 12% gradient BN-PAGE. The bands of Dimer and Monomer after sodium cholate treatment are marked by closed and open triangles. White lead phosphate bands, indicating ATPase activity appeared in the right panel. Molecular masses are shown on the left. (D) Phosphate release assay of Dimer under various ATP concentrations with and without sodium azide. The ATPase activities under 0- and 15.4-mM sodium azide are marked by red and green filled circles, respectively (*n* = 3). These data were fitted by the Michaelis-Menten equation as solid lines.

To examine the assembly of these proteins, we applied the isolated fraction to gel filtration chromatography using a Sephacryl S-400 HR column, which can fractionate up to 8000 kDa globular proteins (Fig. 2B). The proteins eluted as a single peak at a non-void position and were larger than 669 kDa, suggesting that they form a large complex. The molar ratios of the components were estimated to be 3.2:2.9:3.0:1.0:2.3 for MMOBs 1670, 4530, 1660, 1630, and 1620, respectively, from the relative intensity of the SDS-PAGE bands. We then analyzed the isolated fraction by blue-native (BN) PAGE (Fig. 2C, left). A single band was detected, which is consistent with the result of gel filtration chromatography showing a single peak. Next, we applied the band to an In-gel ATPase activity assay, which detects the activity as a white precipitation of lead caused by released of inorganic phosphate (Fig. 2C, right). The band with the complex showed precipitation, whereas the band with thyroglobulin, the negative control, did not. This result indicates that the isolated complex has ATPase activity.

In addition, we assayed the isolated fraction for phosphate release from solution. The complex hydrolyzed ATP at a maximum turnover rate of 0.18 molecules/s per MMOB1670 subunit, β-subunit paralog with a *K_m_* of 74 μM at 25°C (Fig. 2D). The ATPase activity was inhibited by addition of 15.4 mM sodium azide, an inhibitor to ATPases with Walker A motifs (35), with a *K_m_* of 108 μM and a maximum turnover rate of 0.055 molecules/s. In a previous study, the Triton-insoluble fraction, which included the internal structure, showed ATPase activity with a *K_m_* of 66 μ maximum turnover rate of 0.09 molecules/s and was suppressed by 15.4 mM sodium azide, showing a *K_m_* of 84 μM and a maximum rate of 0.063 molecules/s (23). The values obtained here are comparable to these previous data. The above results suggest that the isolated complex is the motor in the internal structure of the gliding machinery. We used this unit complex for further analyses.

### Hexamers resembling F_1_-ATPase catalytic (αβ)_3_ form a dimer

We observed the unit complex by EM using the negative-staining method. A field image showed uniform particles with dimensions of 40 and 20 nm (Fig. 3A and B). As the particle frequency depended on the protein concentration, we concluded that the observed particles were a part of the protein complex. We picked 2148 particle images automatically using RELION software (36) for 2D-classification. From the 2D-classification, we obtained four clear particle images (Fig. S4). We adopted mirror images according to the structure observed in high-speed atomic force microscopy (HS-AFM) (see below). Structural handedness cannot be judged from EM images because they are projections of electrons transmitted through the sample. We focused on an image showing a complex structure with dimensions of 35 and 26 nm featuring nearly two-fold symmetry (Fig. 3C and D). Interestingly, the characteristic hexamer of about 12 nm in diameter formed a dimer and was reminiscent of F_1_-ATPase catalytic (αβ)_3_. Considering that the amino acid sequences of MMOBs 1660 and 1670 have high identity to the α- and β-subunits of F_1_-ATPase, respectively, the dimeric complex is likely an evolutionary related F_1_-ATPase. The distance between the centers of the two hexamers was 11.0 nm. The complex had ten filamentous structures around the two hexamers, four of which appeared to form bridges across the two hexamers. The filamentous structures are unlikely artifact, because they showed common features in four independent averaged images (Fig. S4). Hereafter, we refer to this structural unit as “Dimer.”

**FIG 3.**
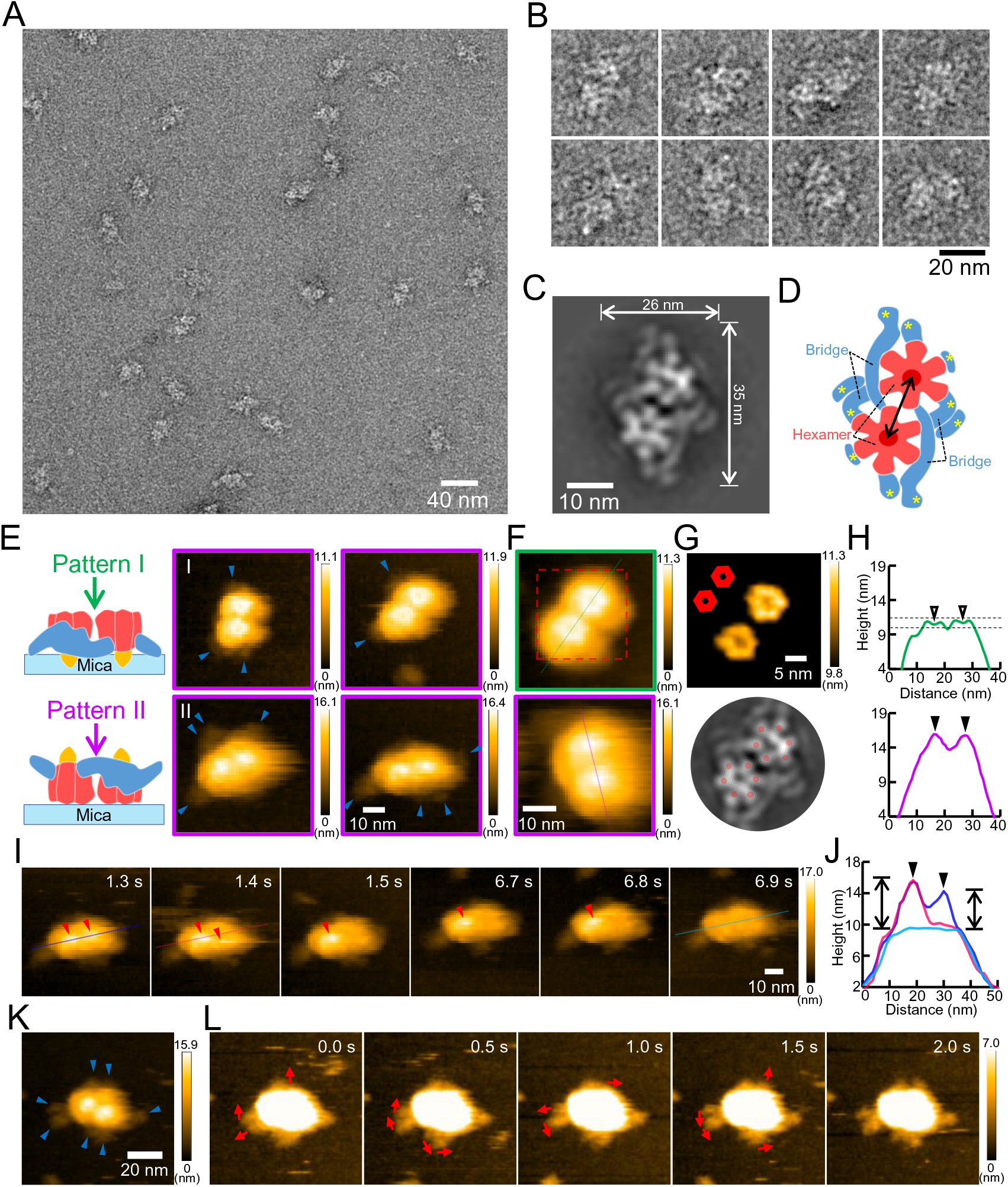
Negative-staining EM and HS-AFM of Dimer. (A) Electron micrograph of negatively stained particles in unit complex fraction. (B) Images of individual particles. (C) Representative 2D averaged image. A mirror image is shown to match the orientation of the hexameric ring observed by HS-AFM. (D) Illustration based on the averaged image in panel C. Filamentous structures are marked by asterisks at an end. The double-headed arrow shows the distance between the centers of the hexamers. (E) Two patterns of HS-AFM images. A Dimer was scanned at 56 × 56 pixels in an area of 70 × 70 nm^2^ with a scanning rate of 100 ms per frame. Illustrations for patterns I and II (left side) were depicted based on 3D chain model shown in Fig. 5. Observation directions are indicated by arrows. Protrusions are marked by blue triangles. Images of patterns I and II are shown in green and purple frames, respectively. (F) Averaged images for patterns I (green frame) and II (purple frame). Dimer was scanned at 50 × 50 pixels in an area of 40 × 40 nm^2^ with a scanning rate of 100 ms per frame. The images were produced by averaging three successive video frames. (G) HS-AFM slice image showing two hexameric rings (upper) and averaged EM image (lower). Upper: The red broken boxed area in panel F was sliced for the height 9.8–11.3 nm from the substrate surface, processed for smoothing, and magnified. The angle alignments of two hexamers are schematically shown in the left upper. Lower: The central part of panel C was excised and aligned to compare with the upper panel. Subunits of the hexamer are marked by red circles. (H) Surface profiles along the lines in pattern I (green) and II (purple). The upper and lower images in panel F were each profiled at the green and purple lines passing the globule centers. The dimples and the peaks are marked by open and black triangles, respectively. The slice height in panel G is shown by broken lines. (I) Shedding process of the peaks of pattern II particle shown in panel E. The peaks are marked by red triangles. (J) Surface profile showing the disappeared peaks. The images in panel I were each profiled at the blue, pink, and light blue lines passing the globule centers. The peaks are marked by black triangles. The double-headed arrows show the peak heights. (K) HS-AFM image of Dimer with seven lateral protrusions. Dimer was scanned at 120 × 120 pixels in an area of 120 × 120 nm^2^ with a scanning rate of 500 ms per frame. Lateral protrusions are indicated by blue triangles. (L) Fluctuations of the protrusions of the particle shown in panel K. The images were sliced for the height 0–7.0 nm from the substrate surface. The moving directions are indicated by arrows. In all HS-AFM images, the color bar on the right shows the range of image heights.

### The hexamer featured a ring and a peak in HS-AFM of Dimer

Next, we visualized Dimer using HS-AFM to clarify the structure under liquid conditions, because the molecules are in dry condition under negative-staining EM. HS-AFM is a powerful method that can visualize the structure and dynamics of single molecules in liquid conditions at a video rate (37, 38). In this method, a specimen is placed on the stage surface and with a probe is scanned in buffer at high speed. In this study, we placed Dimer on a mica surface and scanned it in an area of 70 × 70 nm^2^ at 56 × 56 pixels with a scanning rate of 100 ms per frame. HS-AFM images showed a complex with dimensions of approximately 30 and 20 nm composed of two globules and attached by 2–4 lateral protrusions shorter than 15 nm (Fig. 3E; Movie S1 and S2). The Dimer images were categorized into two patterns as either a ring (pattern I) or a peak (pattern II), based on the central part. Then, we observed them at a higher resolution (area, 40 × 40 nm^2^ with 50 × 50 pixels; scanning rate, 100 ms per frame) (Fig. 3F). In pattern I, the slice image near the top end of the particle between 9.8 and 11.3 nm above the substrate surface showed two hexameric rings (Fig. 3G). The position and direction of the two rings are consistent with those of the hexamers in the negative-staining EM image. In addition, the distance between the centers of the two hexameric rings was 10.4 nm (Fig. 3H), similar with the distance between the centers of the hexamers in the negative-staining EM image (Fig. 3C). These observations suggest that the shape of the Dimer structure in liquid is preserved in negative-staining EM conditions and that the hexamers form rings like F_1_-ATPase catalytic (αβ)_3_. In pattern II, the two central peaks were positioned 11.2 nm apart (Fig. 3H, lower), similar to the distance between the centers of the hexameric rings in pattern I (Fig. 3H, upper), suggesting that patterns I and II are two sides of the same coin (Fig. 3E, left). The distances between the hexameric rings are slightly different for patterns I and II, as 10.4 and 11.2 nm, respectively. This difference suggests that the central axes of two hexamers are not parallel. Considering that HS-AFM detects surface structures while EM shows projection images, the corresponding distance, 11.0 mm in EM is consistent with the numbers obtained from HS-AFM. Interestingly, in most of the particles, the two peaks at 6 and 5 nm became invisible in 20 s, between frames 1 and 3 (Fig. 3I and J; Movie S3). We concluded that these subunits dropped out from the complex, because they did not reappear until the complex was disrupted. Next, we focused on the lateral protrusions of these particles, which may be related to the sheet formation of chains (23). To visualize them more clearly, we scanned Dimer by HS-AFM with an area of 120 × 120 nm^2^, 120 × 120 pixels, and scanning rate 500 ms per frame. Dimer showed seven lateral protrusions around the two globules (Fig. 3K). These protrusions swayed without being fixed (Fig. 3L; Movie S4).

### Isolation and structure of Monomer

To clarify the components and structure of the hexamer, we focused on the monomeric hexamer, called Monomer (Fig. S2). We treated Dimer fraction with 1.5% sodium cholate, an anionic detergent. BN-PAGE and In-gel ATPase activity assays showed a single band with ATPase activity at a position lower than the original one, corresponding to 720–1048 kDa, indicating that Dimer dissociated into smaller units with ATPase activity (Fig. 2C). Therefore, to isolate Monomer, we applied Pellet-1 sequentially to the treatments with 250 mM NaCl for 8 h, 1.5% sodium cholate for 7 h, and to Sephacryl S-400HR gel filtration chromatography (Fig. S2). The elution pattern showed a rather isolated small peak and following overlapping peaks (Fig. S3C). An SDS-PAGE gel showed that MMOBs 1670, 1660, and 1630 eluted in the same fractions, while MMOBs 4530 and 1620 eluted at later fractions (Fig. S3CD), indicating that MMOBs 4530 and 1620 were dissociated from Dimer by sodium cholate treatment. Then, we focused on the small peak fraction, which mainly contained MMOBs 1670, 1660, and 1630 (Fig. 2A, middle, S3D). This complex presumably corresponds to the BN-PAGE band showing ATPase activity (Fig. 2C), because only MMOBs 1660 and 1670 have the Walker A and Walker B motifs in the Dimer components. EM observation using the negative-staining method showed uniform globular particles 10–15 nm in diameter (Fig. 4A and B). As the particle frequency depended on the protein concentration, we concluded that the observed particles were a part of Monomer. We picked 11687 particle images automatically using RELION software for 2D-classification. By 2D-classifing the images in 50 classes, we obtained 15 clear particle images, which were averaged (Fig. S5). Fig. 4C shows a 12 nm diameter globule characterized by a single hexameric ring, corresponding to a part of the Dimer image in Fig. 3C. Three of the subunits were larger than the others with hook structures on either side of the edge. Three averaged images (II–IV) showed a mushroom-like structure resembling F_1_-ATPase, which is characterized by a 12 nm-diameter umbrella and a 3 nm-long stalk (Fig. 4C). Now, we can suggest that MMOBs 1670, 1660, and 1630 form Monomer, i.e., F_1_-like ATPase unit. MMOB4530 was probably not included in this unit because it probably binds to the complex and could not be distinguished in the image due to the low proportion of bound entities (Fig. 2A, middle). Thus, the hexameric ring is likely formed by the α -subunit homolog MMOB1670, and the β-subunit homolog MMOB1670, and the stalk is formed by MMOB1630.

**FIG 4.**
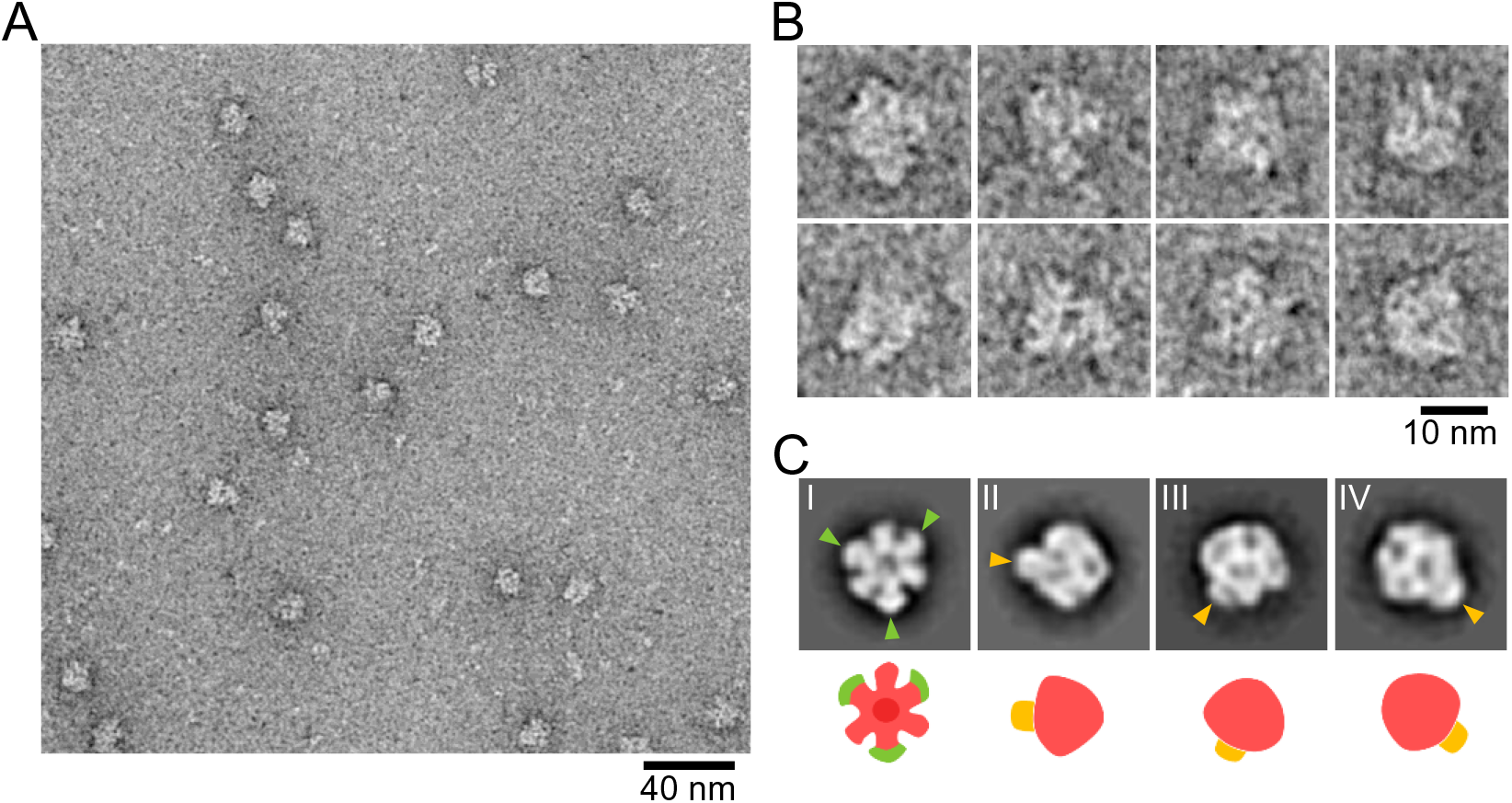
Negative staining EM of Monomer fraction. (A) Electron micrograph of negatively stained ATPase in Monomer fraction. (B) Images of individual particles. (C) Representative 2D averaged images (upper) and depictions of their structures (lower). Upper: Hook structures in the hexameric ring and the stalks are marked by green and orange triangles, respectively. Lower: The hexameric part, hook structures and the stalk are colored rose, green and orange, respectively.

### Isolation and structure of Chain

In gliding machinery, Dimer link to form chains. To characterize these chains, we isolated “Chain fraction” with milder mechanical treatment than those for other fractions. Pellet-1 was treated by 387 mM NaCl and soluble fraction was isolated. Chain fraction contained more than 30 proteins, including the Dimer component proteins MMOBs 1670, 4530, 1660, 1630, and 1620 as major components (Fig. 2A, right). EM observation using the negative-staining method showed chain structures with lengths longer than 70 nm and particles of various sizes (Fig. 5A and B). This time we manually picked 2127 particles from the chain images, overlapping approximately 50% of the 71 × 71 nm^2^ box area. From 2D-classifcation, we obtained seven clear particle images (Fig. 5C). The particle images show the various orientations required for 3D reconstruction. We then created a 3D map by combining a total of 1709 particle images of good quality (Fig. 5D and S6). The 3D map with dimensions of 70, 20, and 15 nm at a density threshold (contour level = 0.026) was composed of two dimers of mushroom-like structures resembling F_1_-ATPase, aligned along the dimer axis (Fig. 5D). Dimers were connected by a bulge structure with a diameter of 5 nm. The chain interval was 31 nm, consistent with the corresponding dimension in a 2D image from electron cryotomography (ECT) (23), suggesting that the 3D model obtained reflects the original structure from a cell. The mushroom-like structure with a diameter of 15 nm, consisting of a hexameric ring and a central stalk, was connected to the dimer by two bridge structures with a diameter of 3–6 nm. An atomic model of F_1_-ATPase catalytic (αβ)_3_ from *Bacillus* PS3 (PDB ID 6N2Y) (39) was fitted into each hexameric ring of the mushroom-like structure (Fig. 5E). The distance between the centers of the fitted (αβ)_3_ in Dimer was 12.5 nm, which is consistent with that of Dimer observed by negative-staining EM and HS-AFM (Fig. 3C and H). The fitted model showed that each hexameric ring had two protrusions of 3–6 nm pointing laterally (Fig. 5E). The cross-sections of each mushroom-like structure showed the central stalk length of 5 nm (Fig. 5F). A cavity was observed at the center of the hexameric ring. However, it may be an artifact of the low-resolution map of negative-staining EM, because metal coating tends to emphasize the peripheral part of large particles (40). Next, we compared a reprojection image of the 3D chain map with the 2D averaged image of Dimer from negative-staining EM (Fig. 5G). Two short filaments marked by asterisks in Dimer (Fig. 5G, left) are positioned facing each other in the connecting bulge (Fig. 5G). Previously, ECT of a permeabilized *M. mobile* cell showed a chain structure characterized by repeats of two globules and two types of projections to the cell membrane (Fig. 5H, left) (23). The hexameric ring and the central stalk in the 3D map here correspond to the globule and one type of projection to the cell membrane in the ECT image, respectively, suggesting that the chain is oriented with the central stalk facing the membrane, which is common in F_1_-ATPases. At the interface between Dimers, the 3D map here did not include a structure composed of another type of projection and a globule as observed in the ECT image. The subunits corresponding to these structures probably had structural variations or dissociation during the isolation process.

**FIG 5.**
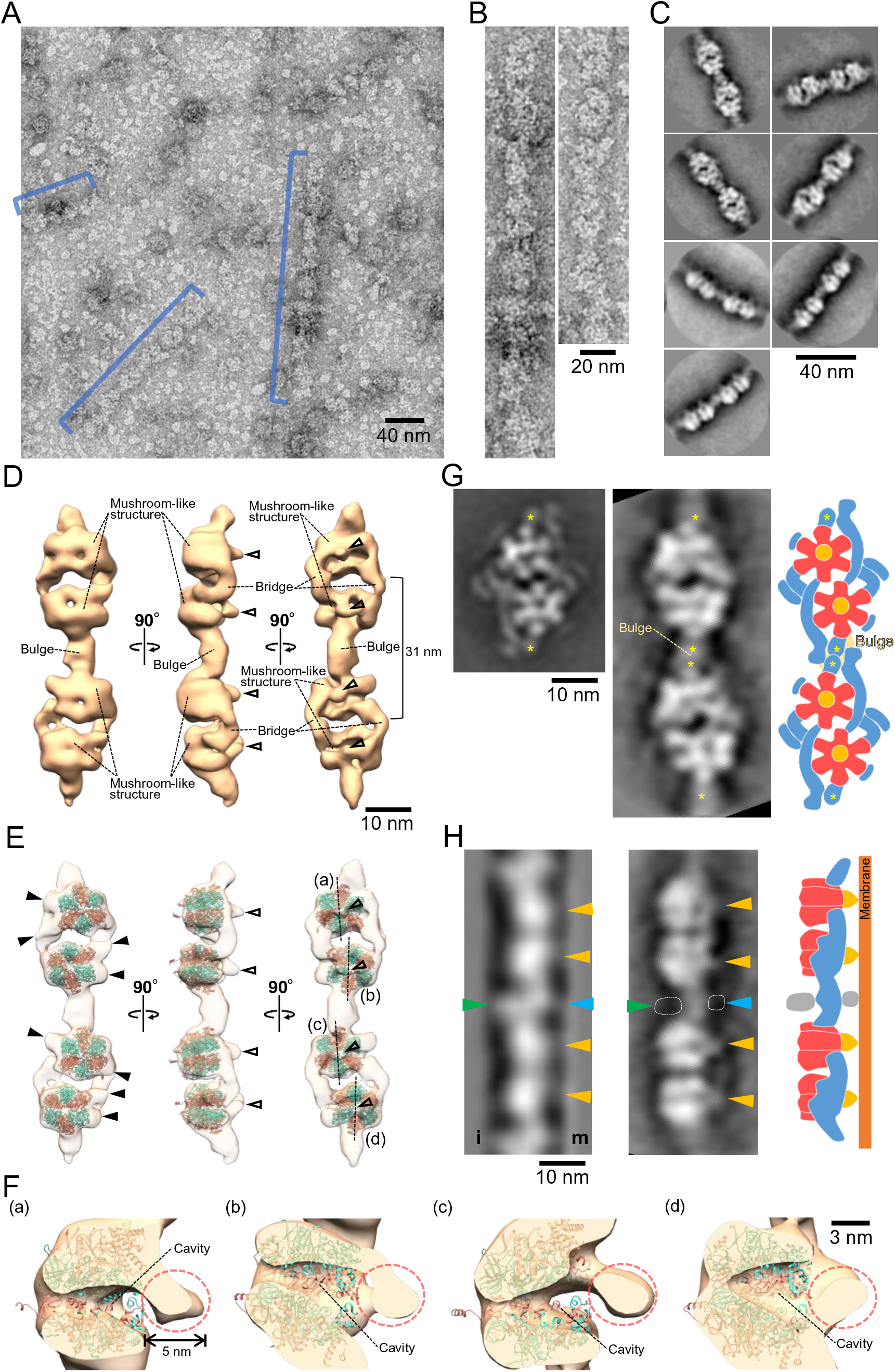
Chain structure. (A) Electron micrograph of negatively stained Chain. The chain structures are marked by blue lines. (B) Magnified chain images. (C) Representative 2D averaged images. (D) Three-dimensional reconstruction of Chain containing two F_1_-like ATPase dimers. The 3D map is visualized at a density threshold (contour level = 0.026). The central stalks are marked by open triangles. (E) Superposition of the atomic model of *Bacillus* F_1_-ATPase catalytic (αβ)_3_ (PDB ID 6N2Y) (39) onto the 3D chain structure. The α and β subunits are colored salmon and turquoise, respectively. The central stalk and protrusions from hexameric rings are marked by open and black triangles, respectively. (F) Cross section of mushroom-like structures. Central stalks are marked by broken circles. The double-headed arrow shows the length of the protrusion. Corresponding mushroom-like structures are marked (a)–(d) in panel E. (G) Comparison between Dimer image from Fig. 3C (left) and the Chain reprojection (middle). The reprojected image is viewed from the angle used for the right image of panel D. Short filaments corresponding to the position of the connecting bulge are marked by asterisks. A depiction of Chain model based on the comparison (right). (H) Comparison between the averaged chain image from ECT (left) and the chain reprojection (middle). Left: The averaged chain image was modified from the whole cell ECT image in the previous study (23). Middle: Chain was reprojected from an angle close to the middle image in panel D. Inner sides and membrane relative to the chain are marked by i and m, respectively. The protrusion from the globule corresponding to the central stalk from the hexameric ring, one from the connecting bulge and the globule attached to the connecting bulge are marked by orange, light blue and green triangles, respectively. The areas of image densities that were visualized only in the ECT image are marked by broken lines. Right: An illustration depicts a chain model based on the comparison.

## Discussion

### Outline of internal structure of gliding machinery

Previously, sequence analysis suggested that the chain of *M. mobile* gliding machinery evolved from F_1_-ATPase (5, 23–26). The present study supports this conclusion by structural data showing that the chain has hexameric rings similar to the F_1_-ATPase catalytic (αβ)_3_. Integrating available information, we can now describe the outline of the internal structure of *the M. mobile* gliding machinery (Fig. 1A). *M. mobile* cells have 28 individual 530 nm long chains, each of which contains 17 Dimer units composed of two F_1_-like ATPases and filamentous structures (23). The central stalk of the F_1_-like ATPase and another protrusion from the connecting bulge project to the cell membrane.

### Unique role of F_1_-ATPase related complex

To date, several complexes are known to be evolutionarily related to F_1_-ATPase, all of which are responsible for transporting substances across the membrane (41). However, the motor we identified here most likely plays a role in motility. This case may be reminiscent of dynein, a motor in eukaryotes, which evolved from a widely conserved AAA (ATPases associated with diverse cellular activities)+ protein, in which multiple subunits of ATPases perform functional rotation (42, 43). Sequence analyses have shown that mycoplasma Type 3 ATPase is also related to F_1_-ATPase, and its role has been suggested to promote substrate turnover in the MIB-MIP system (33). If Type 3 ATPase provides the force to change the conformation of a hydrolytic enzyme, its role in force generation is common with Type 2, the gliding motor.

F_1_F_o_-ATPases are known to be dimerized through interactions between F_o_-domains and are usually arranged in rows along the short axis in the tightly curved cristae ridges of mitochondria (31, 44, 45). The dimer structure found in the present study is not related to this, because the F_1_-like domain is dimerized through the filament structure and is linked in the long axis direction. However, some roles may be common in part if the dimerization and chain formation observed in the gliding motor identified in this study stabilizes the membrane structure, as seen in the F_1_F_o_-ATPase dimer (46). Moreover, dimerization may result in cooperativity in motor functions.

### Protein assignment

The α-subunit homolog MMOB1660 (58.7 kDa) and the -subunit homolog MMOB1660 (58.7 kDa) and the β-subunit homolog MMOB1670 (88.4 kDa) likely correspond to the smaller and larger subunits, respectively, of the hexameric ring of an F_1_-like ATPase unit, as suggested by the estimated 1:1 molar ratio in Dimer (Fig. 2A, 4C). This means that the hook structure of the larger subunit may be formed by the extra N-terminal region (34.8 kDa) of MMOB1670. Previously, 3D structure modeling based on secondary structure suggested that MMOB1630 is structurally similar to the γ subunit, the principal component of the central stalk of F_1_-ATPase (5). In general, the γ subunit of F_1_-ATPase is composed of a coiled-coil and a globular domain and penetrates the hexameric ring (47). In the F_1_-like ATPase unit and chain 3D model, a stalk structure, suggesting the globular domain of the γ subunit, was found in the center of the hexameric ring (Fig. 4C and 5F), implying that MMOB1630 penetrates the hexameric ring like the γ subunit.

Using HS-AFM observations, the peak at approximately 5 nm at the center of the hexameric ring dropped out with time (Fig. 3I). The peak height agrees with the length of the estimated globular domain of MMOB1630 in Chain 3D model (Fig. 3J and 5F), suggesting that the peak is composed of MMOB1630 and was pulled out from the hexameric ring by the scanning cantilever during HS-AFM observation. The pull-out event is thought to be common to that of the F_1_-ATPase, in which the γ subunit is removed from the hexameric ring by optical tweezers (48).The filamentous structures around the hexameric ring probably correspond to lateral protrusions in the HS-AFM images and are formed by the remaining proteins, MMOB1620 and MMOB4530 (PGK). These proteins probably play roles in ATPase dimerization, chain formation, and lateral chain interaction (23). MMOB1620 is an unannotated protein specific to the Type 2 ATPase cluster (26, 32). MMOB4530 is annotated as an enzyme that transfers phosphate groups from 1,3-bisphosphoglycerate to ADP in glycolysis to yield ATP and 3-phosphoglycerate (49). In *M. mobile*, ATP is probably provided by glycolysis (50). MMOB4530 may supply ATP efficiently to the gliding motor by its close proximity. Yeast V-ATPase, which belongs to the rotary ATPase family like F_1_F_o_-ATPase, is also attached by two glycolytic enzymes, 6-phosphofructo-1-kinase and aldolase (51–53). These glycolytic enzymes are involved in the regulation of V-ATPase assembly and activity.

Ten proteins have been identified as the components of the internal structure (Fig. 1) (23, 24, 26). In the present study, we identified five proteins as the Dimer components. Other five proteins are likely involved in other parts, for example “bell” at the front part of internal structure (Fig. 1A) and parts of the intact chain structure which were lost in the fractionation (Fig. 5H).

### Possible force transmission mechanisms for gliding

The involvement of an internal ATPase in the gliding mechanism is based on the following five observations from the analysis of the “gliding head” of *M. mobile* protrusions and of the isolated gliding machinery: (a) The affinity for ATP estimated by the saturation extent is comparable in the ATPase activity of the internal structure and the speed of the gliding head (23). (b) Substrate binding and gliding speed of the gliding head are inhibited by azide, as well as the ATPase activity of the internal structure (23). (c) The chain in the internal structure undergoes conformational changes based on ATP hydrolysis (23, 34). (d) Among the 21 proteins identified from the gliding head, only MMOBs 1660 and 1670 could be suggested for ATPase from the amino acid sequences alone (23, 24). (e) Fluorescent protein tagging of components of the internal structure significantly affects the substrate binding of cell and the gliding speed (26).

The structure elucidated in the present study allows us to discuss the gliding mechanism in more detail. In F_1_-ATPase, the three catalytic sites in the hexameric ring cooperatively hydrolyze ATP, and each catalytic β-subunit undergoes a conformational change that drives the rotation of the central stalk (47). Previously, structural changes linked to ATP hydrolysis were reported: (I) EM studies showed 2 nm contraction of Dimer intervals in the isolated chains (23), and (II) HS-AFM studies showed movements of individual Dimers in the cell 9 nm perpendicular to the chain long axis and 2 nm into the cell (34). Considering these observations, we propose two different working models for the force transmission mechanism in gliding (Fig. 6). In the “contraction model” (Fig. 6 i), the force generated by the hexameric ring shortens the chain. The resulting displacement of the projections from Dimer to the cell membrane drives the hook structure of Gli521 like a “lever.” Then, the leg moves with the catch, pull, and release of the SOs. In the “rotation model” (Fig. 6 ii), the force generated by the hexameric ring rotates MMOB1630 in the same way as F_1_-ATPase. This rotation is transmitted across the cell membrane to the Gli521. The hook structure converts rotational motion into linear motion of the leg, similar to a crank. Previous studies have reported that *M. mobile* exhibits unitary steps of approximately 70 nm in size at no load (29, 54). In our models, both the rotation and contraction displacements are expected to be a few nanometers. These displacements may be amplified by the large surface structure complex formed by the 100 nm long Gli349 and 120 nm long Gli521, which show dimensions comparable to the step size, acting as a large gear (22). This conjecture could explain how a single leg exerts a force of 1.5 pN, smaller than that of conventional motor proteins such as myosin, dynein, and kinesin.

**FIG 6.**
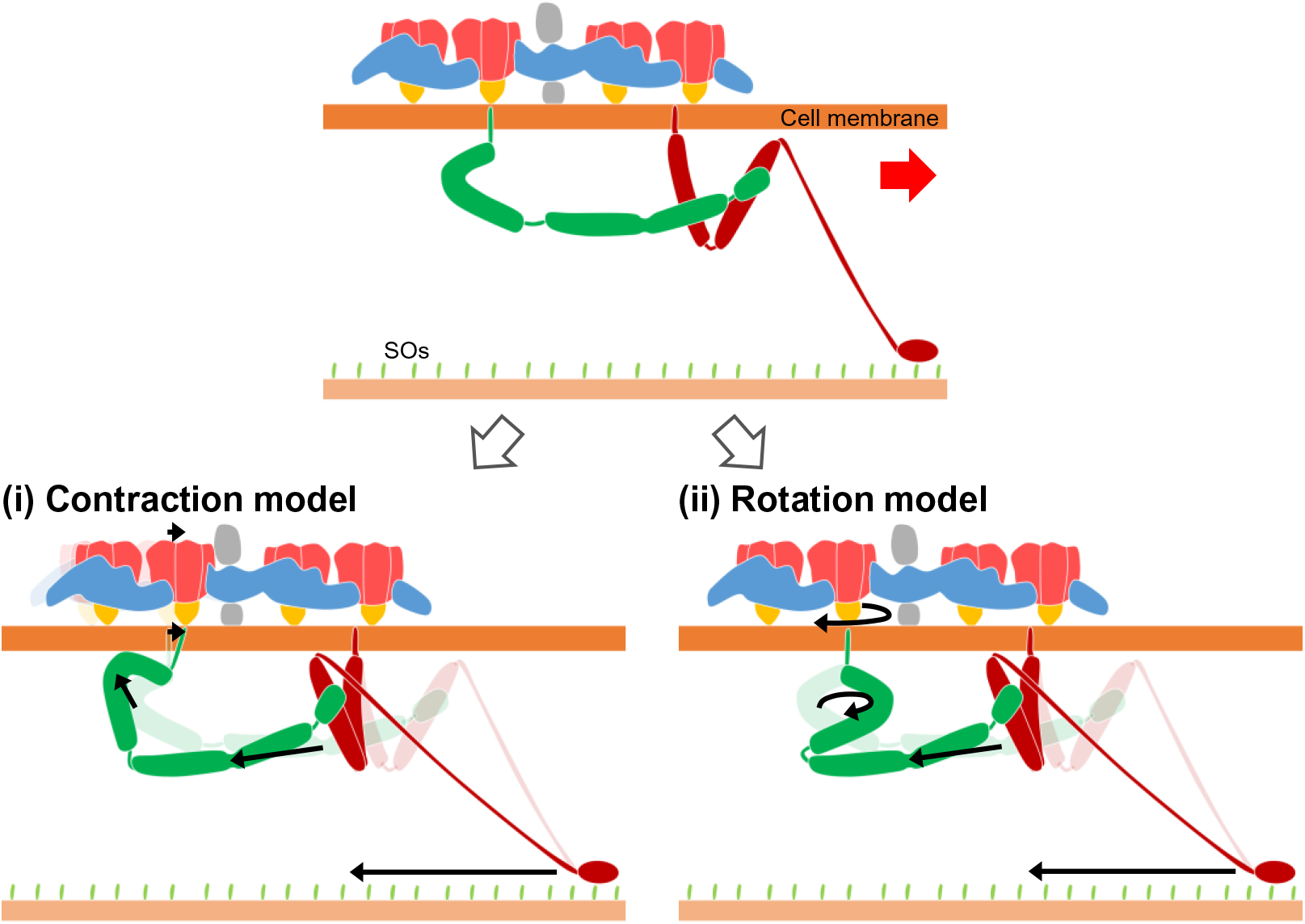
Working models for force transmission mechanism. The gliding direction is indicated by a red arrow. The regions marked in gray were visualized only in the ECT image. The crank protein Gli521 and the leg protein Gli349 are colored green and red, respectively. (i) Contraction model: The force generated by the hexameric ring displaces Dimer along the gliding direction. The displacements are transmitted through the membrane to Gli521. (ii) Rotation model: The force generated by the hexameric ring rotates the central stalk in a mechanism similar to that of F_1_-ATPase. The rotational motion is transmitted across the membrane to Gli521.The generation and transmission of forces are presented by black arrows for both models.

### Evolution of *M. mobile* gliding

A previous study suggested that Gli349 evolved from a static binding receptor to parasitize the host (16). Considering this, the evolutionary origin of *M. mobile* gliding can now be discussed. F_1_F_o_-ATPase, which is abundant on the cell membrane, could have been accidentally associated with the binding receptor and turned into a primitive motility system, which may have provided random cell spreading. The system was then refined under survival pressure, because motility might be beneficial to infection and evading the host’s immune system. For dimerization and chain formation, PGK was then incorporated into the gliding machinery, because PGK was working in close proximity to F_1_F_o_-ATPase.

## Materials and Methods

### Strains and culture conditions

We used P476R *gli521*, a mutant strain of *M. mobile* that can glide normally but binds SOs more tightly than wild-type strains (10, 22, 28). *M. mobile* cells were cultured as described previously (55, 56).

### Optical microscopy

The cultured cells were inserted into a tunnel chamber assembled with two coverslips and double-sided tapes and observed by phase-contrast microscopy using an inverted microscope (IX71; Olympus, Tokyo, Japan) (17, 19). Movement was recorded using a complementary metal-oxide semiconductor (CMOS) camera (DMK33UX174; The Imaging Source, Bremen, Germany). Video was analyzed using the ImageJ software, version 1.53a (http://rsb.info.nih.gov/ij/).

### Solubility test

All procedures for fractionation were performed at 4°C unless otherwise noted and focused protein bands were identified by peptide mass fingerprinting (PMF), as previously reported (24, 57). To investigate the solubility of the chain components, *M. mobile* cells from 60 mL of culture medium were collected by centrifugation at 14000 × *g* for 20 min and washed twice with PBS consisting of 8.1 mM Na_2_HPO_4_, 1.5 mM KH_2_PO_4_, pH 7.3, 2.7 mM KCl and 137 mM NaCl. Cells were resuspended in PBS to a 12-fold higher concentration than the culture and sonicated for 1 min at 24–27°C to be dispersed in microtubes using an ultrasonic generator (2510 J-MT; BRANSON, Kanagawa, Japan). The cells were then treated with Triton solution (1% Triton X-100, 0.1 mg/mL DNase, 5 mM MgCl_2_, and 1 mM phenylmethylsulfonyl fluoride in PBS) in a total volume of 10 mL. After gentle shaking for 1 h, the suspensions were centrifuged at 20000 × *g* for 20 min, and pellets were collected and washed once with suspension buffer (PBS including 5 mM MgCl_2_). This fraction is Pellet-1. Pellet-1 was then resuspended separately in suspension buffer adjusted to contain different concentrations of NaCl (0, 50, 137, 200, and 400 mM) by pipetting several times. After treatment for 8 h, suspensions were centrifuged at 20000 × *g* for 20 min, and supernatants and pellets collected for SDS-PAGE analysis.

### Preparation of fractions

For isolation of Dimer, Pellet-1 from 1.2-liter culture was resuspended in 5 mL suspension buffer by pipetting up and down and allowed to dissolve for 8 h. The soluble fraction was collected by centrifugation at 20000 × *g* for 20 min and loaded onto a HiLoad 16/600 Superdex 200 column (Cytiva, Tokyo, Japan) equilibrated with 1 mM MgCl_2_ in PBS at a flow rate of 0.8 mL/min. The fractions were analyzed by SDS-PAGE and CBB staining.

For isolation of Monomer, Pellet-1 from 1.2-liter culture was suspended by 5 mL Tris buffer consisting of 20 mM Tris-HCl (pH 7.5), 250 mM NaCl, 1 mM phenylmethylsulfonyl fluoride, and 1 mM MgCl_2_ with pipetting and allowed to dissolve for 8 h. The soluble fraction was collected by centrifugation at 20000 × *g* for 20 min and mixed with 1.5% (w/v) sodium cholate. After 7 h of incubation, the complexes were loaded onto a Sephacryl S-400 HR column (Cytiva) equilibrated with 0.7% sodium cholate, 20 mM Tris-HCl (pH 7.5), 250 mM NaCl, and 1 mM MgCl_2_ at a flow rate of 0.5 mL/min. The fractions were analyzed by SDS-PAGE and CBB- and reverse-staining (58, 59). The fraction of the complex composed of MMOBs 1670, 1660, and 1630 was collected. Samples were concentrated using an Amicon Ultra 100 K spin filter (Merck KGaA, Darmstadt, Germany), if necessary.

For isolation of Chain, Pellet-1 from 15 mL culture was resuspended in 60 μL suspension buffer. The suspension was then gently mixed with an equal volume in a suspension buffer adjusted to contain 637 mM NaCl. Chain was recovered as the supernatant by centrifugation at 5000 × *g* for 5 min.

### Analytical gel filtration

Dimer solution was loaded onto a Sephacryl S-400 HR column equilibrated with gel filtration buffer containing 20 mM Tris-HCl (pH 7.5), 200 mM NaCl and 1 mM MgCl_2_ at a flow rate of 0.5 mL/min at 4°C. Thyroglobulin (669 kDa; Gel Filtration Calibration Kits; Cytiva) was dissolved in gel filtration buffer and loaded onto the column as a size standard at a flow rate of 0.5 mL/min. The stoichiometry of protein complexes was estimated by densitometry of SDS-PAGE gels stained with CBB, using a scanner (GT-9800F; Epson, Nagano, Japan) and ImageJ (9).

### BN-PAGE and in-gel ATPase activity assays

BN-PAGE was performed according to the user manual of the Native PAGE Novex Bis-Tris Gel System (Thermo Fisher Scientific, Waltham, MA). For BN-PAGE of sodium cholate treated Dimer, SC treated, Dimer fraction was mixed with sodium cholate (1.5%) and incubated at 4°C for 9 h. When this sample was mixed with a sample buffer, NativePAGE™ 5% (w/v) G-250 sample additive was also added at 0.5% (w/v) to prevent protein aggregation. Thyroglobulin was dissolved in water and used as a negative control for the In-gel ATPase activity assay. For the In-gel ATPase activity assay (60, 61), native gels were incubated with gentle shaking for 8h at 24–27°C in activity buffer containing 270 mM glycine, 35 mM Tris (pH 8.4), 4 mM ATP, 14 mM MgSO_4_, and 0.2% (w/v) Pb(NO_3_)_2_. The gels were rinsed once with water and images were taken using ImageQuant LAS 4000 mini (Cytiva). White precipitates were then dissolved by gentle shaking for 8 h at 24–27°C with 50% (v/v) methanol and 10% (v/v) acetic acid in water. The gels were restained with 0.025% (w/v) CBB G-250 and 10% acetic acid in water for 80 min at 24–27°C with gentle shaking and destained with 10% (v/v) ethanol and 10% acetic acid in water for 180 min at 24–27°C with gentle shaking. The gels were rinsed once with water and images were taken using ImageQuant LAS 4000 mini.

### Phosphate-release assay

Dimer solution was dialyzed for 8 h using 20 mM Tris-HCl (pH 7.5), 150 mM NaCl, and 2 mM MgCl_2_. ATPase activity was assayed by a continuous spectrophotometric method using a 2-amino-6-mercapto-7-methylpurine ribonucleoside–purine nucleoside phosphorylase reaction to detect released inorganic phosphate (EnzChek kit; Thermo Fisher Scientific) (62). The reaction mixture was as follows: 15.7 nM Dimer, 20□mM Tris-HCl (pH 7.5), 150 mM NaCl, 2□mM MgCl_2_ and 0.01–1 mM ATP in a total volume of 0.2□mL at 25°C. Sodium azide was added to 15.4 mM final concentration when the reaction was started. The protein amount of the MMOB1670 comprising F_1_-ATPase β-subunit paralogs was estimated using densitometric of SDS-PAGE.

### Negative-staining EM and image processing

Dimer solution was placed on a glow-discharged (PIB-10; VACUUM DEVICE, Ibaraki, Japan) carbon-coated grid (F-400; Nisshin EM Co., Tokyo, Japan) and incubated for 1 min at 24–27°C. The solution was then removed, and the grid was stained with 2% uranyl acetate (w/v) for 30 s. The stain was then removed, and the grid was air-dried. To observe Monomer and Chain, the grids were washed with water after 1 min of incubation and then treated as described for Dimer solution. Samples were observed using a transmission EM (JEM1010, JEOL) at 80 kV, equipped with a FastScan-F214 (T) charge-coupled-device (CCD) camera (TVIPS, Gauting, Germany), and images were captured at 2.58 Å/pix.

The contrast transfer function parameters for electron micrographs were estimated using Gctf (63). Further image processing was performed using RELION 3.0 (36). A total of 2148 particles for Dimer and 11687 particles for Monomer were automatically selected with box sizes of 180 × 180 and 100 × 100 pixels using reference-based auto-picking, respectively. These particle images were binned to 5.16 Å/pix. For Dimer, the particle images were 2D-classified into 100 classes. For Monomer, particle images were 2D-classified in four rounds, and the selected 7381 particles were re-extracted with the pixel size returned to the unbinned image and then 2D-classified into 50 classes.

For reconstruction of the 3D chain structure, 2127 particles were manually selected for chains with a box size of 276 × 276 pixels with ∼50% overlap. These particle images were binned to 5.16 Å/pix. Particle images were 2D-classified in two rounds, and the selected 1709 particles were used to reconstruct the initial model with a final resolution limit of 50 Å. The initial model and selected particles were used to perform 3D refinement. Reprojection images were produced from the 3D map using the relion_project command in RELION. The 3D map was visualized using UCSF Chimera 1.14 (64). Atomic models of F_1_-ATPase catalytic (αβ)_3_ from *B. subtilis* (PDB ID 6N2Y) (39) were fitted into the 3D map using command of “Fit in map” in UCSF Chimera.

### HS-AFM

Imaging was performed with a laboratory-built tapping mode HS-AFM (65, 66), using small cantilevers (BLAC10DS-A2, Olympus; resonant frequency, ∼0.5 MHz in water; quality factor, ∼1.5 in water; spring constant, ∼0.1 N/m). The cantilever’s free-oscillation peak-to-peak amplitude (*A*_0_) and set-point amplitude were set at ∼2.5 nm and ∼0.8 × *A*_0_, respectively. The probe tip was grown on the original tip end of a cantilever through electron beam deposition and further sharpened using a radio frequency plasma etcher (PE-2000, South Bay Technology, Redondo Beach, CA) under an argon gas atmosphere (typically at 180 mTorr and 15 W for 3 min). The sample was deposited on a freshly cleaved mica disc glued to a glass stage beforehand. After 3.5 min, the stage surface was immersed in a liquid cell containing an observation buffer [20 mM Tris-HCl (pH 7.5), 50 mM KCl, 2 mM MgCl_2_]. Imaging was performed at 24–27°C. AFM images were processed using a low-pass filter to remove spike noise and make the *xy*-plane flat and analyzed using Kodec software (version 4.4.7.39) (67). Surface profiles and smoothing were performed using ImageJ software.

## Acknowledgments

We thank Toshiaki Arata, Ikuko Fujiwara, Kohei Kobayashi, and Hiroki Sato at the Graduate School of Science, Osaka City University, and Takayuki Uchihashi at the Department of Physics and Structural Biology Research Center, Nagoya University, for helpful discussions. We also thank Aya Takamori at the Graduate School of Science, Osaka City University, for performing MALDI-TOF MASS spectrometry. TT learned RELION software in an instruction course on 20180927–28 provided by the Cyclic Innovation for Clinical Empowerment (CiCLE) from AMED.

This work was supported by a Grant-in-Aid for Scientific Research on the Innovative Area “Harmonized Supramolecular Motility Machinery and Its Diversity” (MEXT KAKENHI Grant Number JP24117002), by a Grants-in-Aid for Scientific Research (B) and (A) (MEXT KAKENHI Grant Numbers JP24390107, JP17H01544), by JST CREST Grant Number JPMJCR19S5, Japan, by the Osaka City University (OCU) Strategic Research Grant 2018 for top priority researches and by a Grant-in-aid of the FUGAKU TRUST FOR MEDICINAL RESEARCH to MM, and JSPS KAKENHI (Grant Number JP25000013), the Platform Project for Supporting Drug Discovery and Life Science Research (BINDS) from AMED (Grant Number JP19am0101117 and support number 1282), CiCLE (Grant Number JP17pc0101020), and JEOL YOKOGUSHI Research Alliance Laboratories of Osaka University to KN.

## Figure Legends

**FIG S1.**
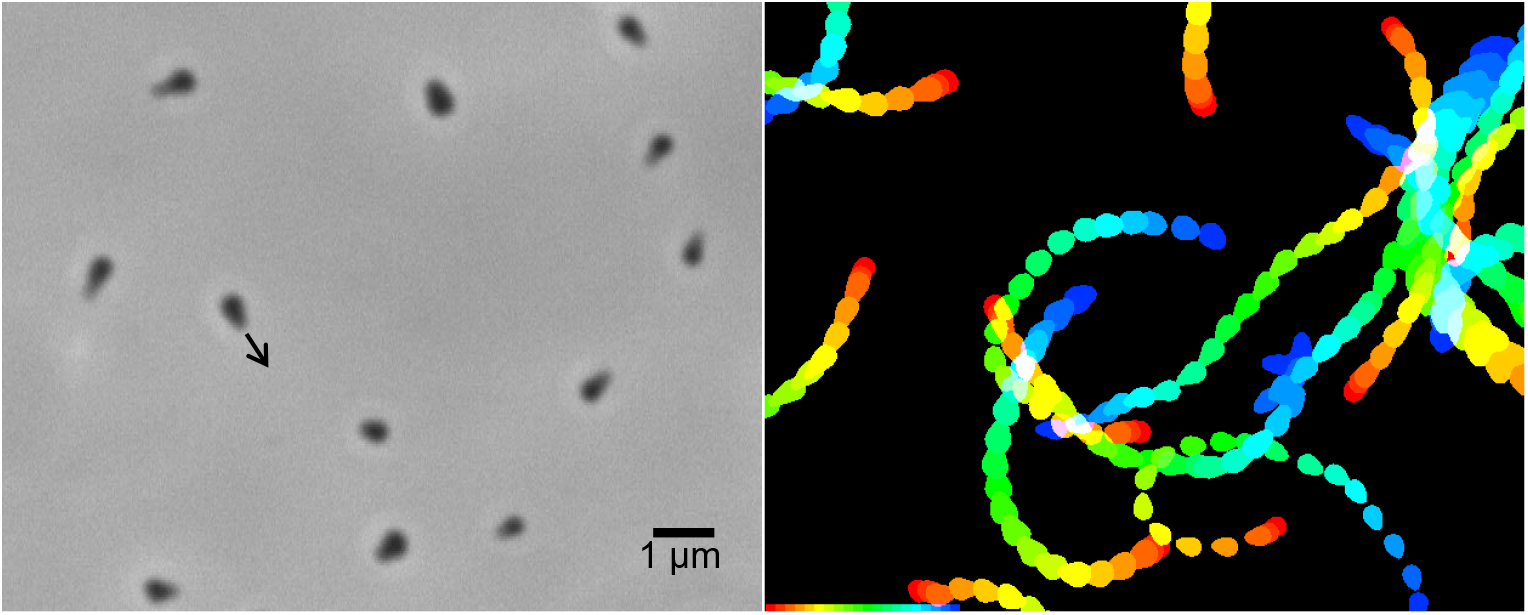
Gliding of *M. mobile* cells. Optical microscopy of cells (left) and trajectories of gliding cells (right). All cells are gliding in the direction of tapered end as indicated by a black arrow. For trajectories, video frames of every 0.2 s were colored differently from red to blue, and stacked for 4 s.

**FIG S2.**
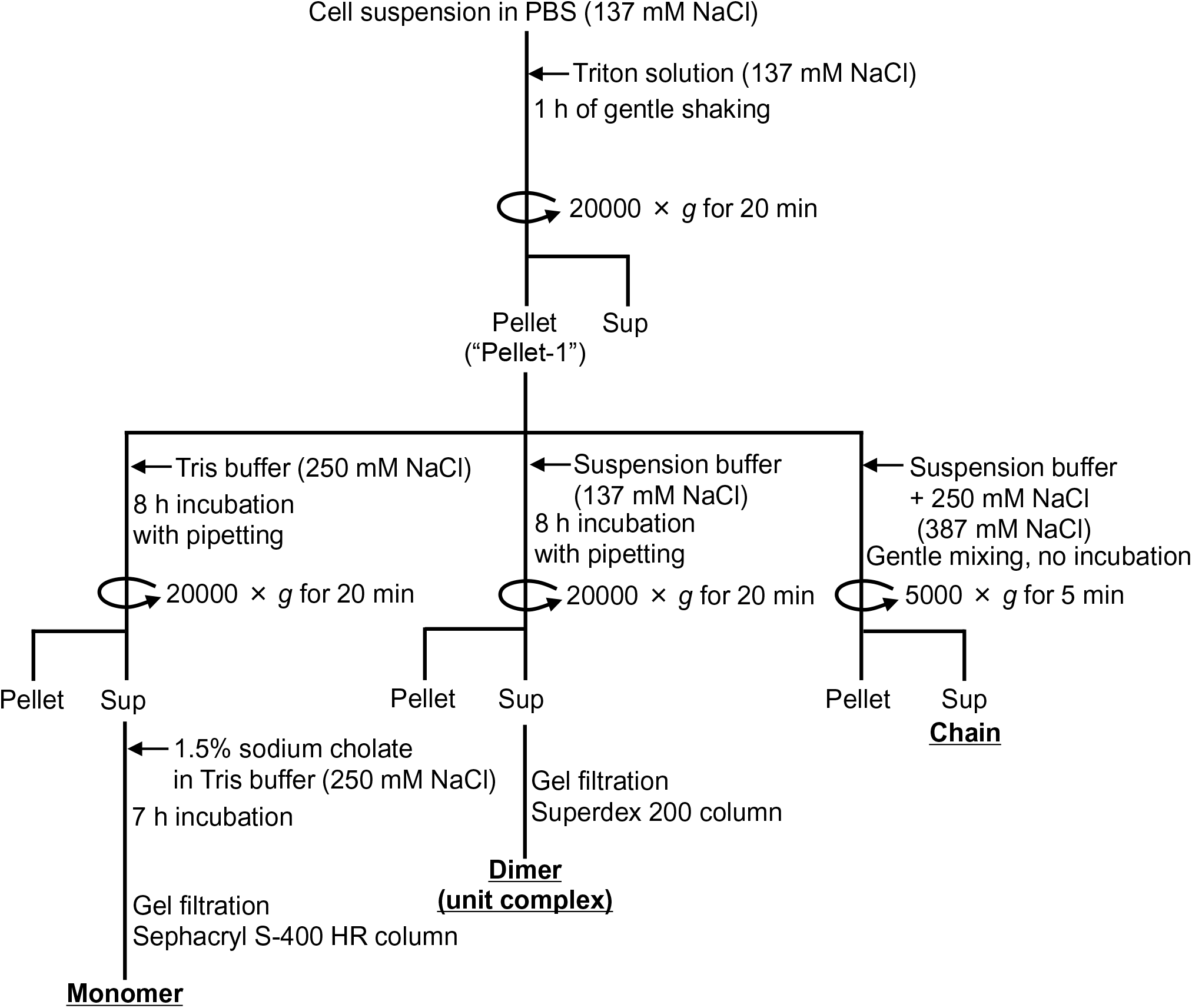
Isolation procedure for three fractions. Each fraction was obtained from Pellet-1 fraction. The adjusted concentrations of NaCl in the solution are indicated in parentheses. Supernatant is abbreviated as “Sup”. See Materials and Methods for details.

**FIG S3.**
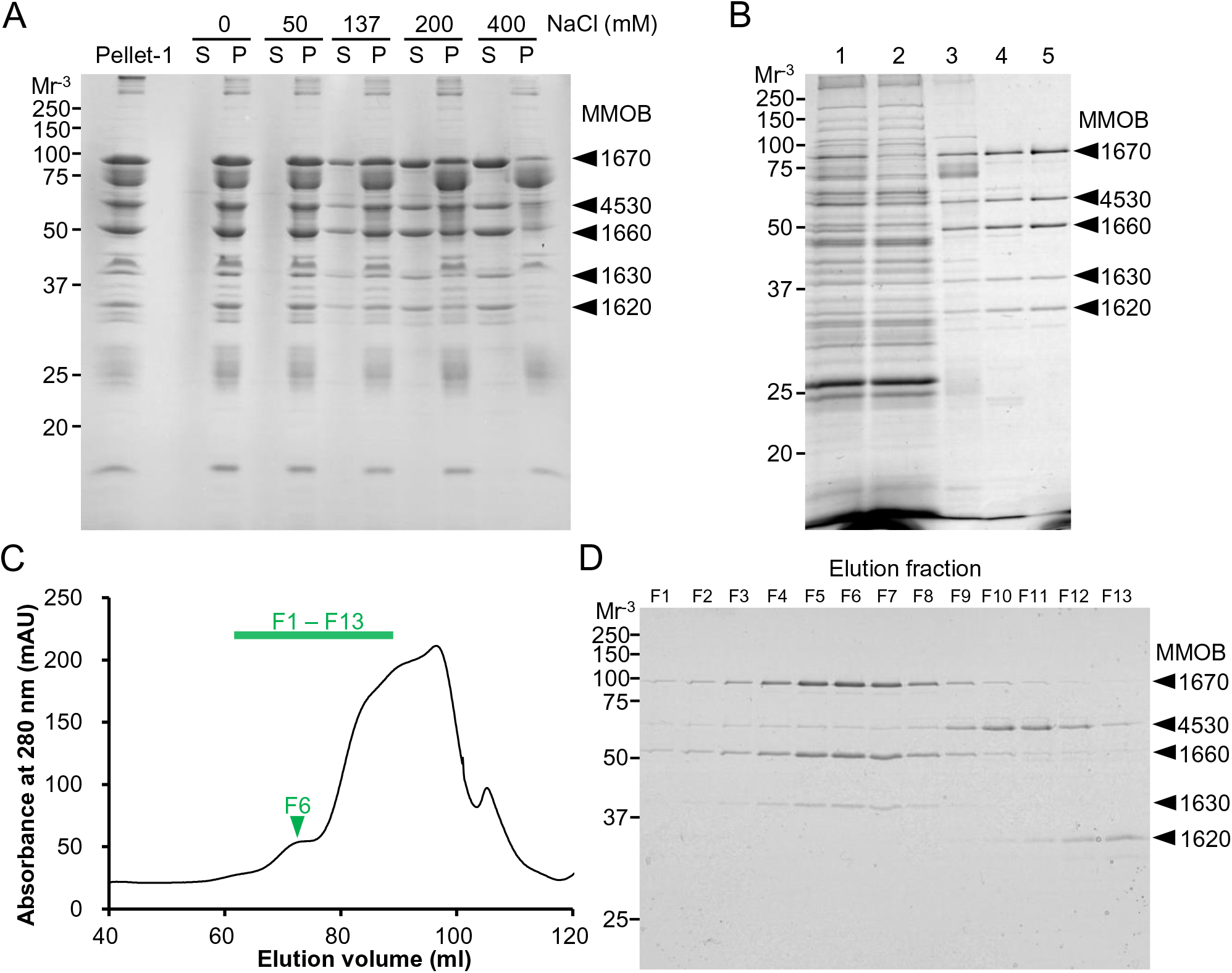
Isolation profile of Dimer (unit complex) and Monomer fractions. (A) Solubility of the chain components. Pellet-1 fraction was treated with buffers containing the specified concentrations of NaCl and centrifuged. Pellet-1 fraction, the supernatants (S) and the pellets (P) were analyzed by SDS-12.5% PAGE. See Materials and Methods for details. (B) Isolation of Dimer (unit complex). The fractions of each step were subjected to SDS-12.5% PAGE gel and stained with CBB. Lane 1, lysate of *M. mobile* cells; lane 2, Triton-soluble fraction; lane 3, Triton-insoluble (Pellet-1) fraction; lane 4, supernatant after incubation in suspension buffer containing 137 mM NaCl; lane 5, peak fraction of Superdex 200 gel filtration chromatography. The bands of Dimer components are marked by black triangles. (C) Gel filtration profile in Monomer isolation using a Sephacryl S-400 HR column. The small peak is marked by a green triangle. (D) The F1–F13 fractions in the gel filtration indicated by the green line in panel C were subjected to SDS-12.5% PAGE and stained with CBB. F6, which corresponds to the small peak in the gel filtration, was used for further analyses as Monomer fraction. Bands of Dimer components are marked by black triangles. For panels (A, B, D), Focused proteins were identified by PMF, and molecular masses are shown on the left.

**FIG S4.**
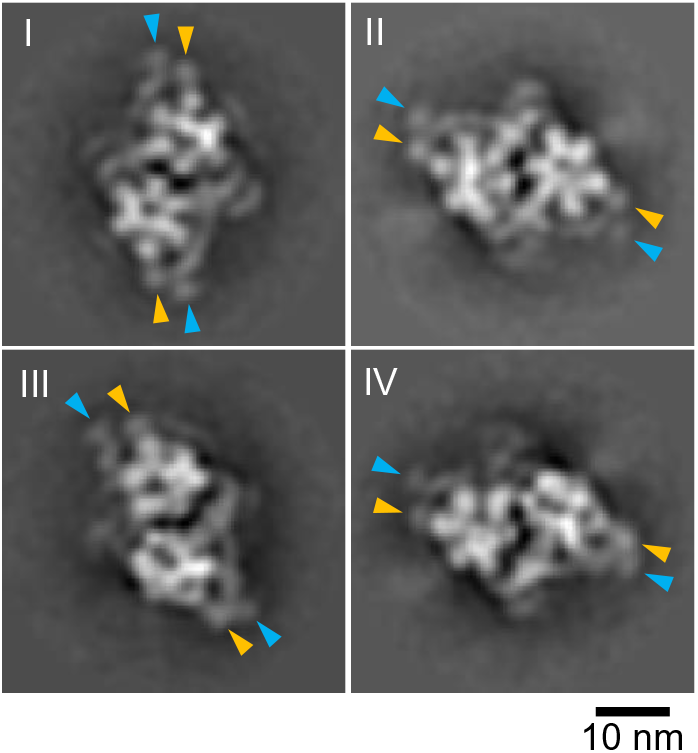
Two-dimensional averaged images of Dimer obtained by negative-staining EM. Four classes of clear particle images from 20 classes are represented. These images have common features as represented long and short protrusions marked by light blue and orange triangles, respectively. As mentioned in Fig. 3C, the images were mirrored to uniform handedness.

**FIG S5.**
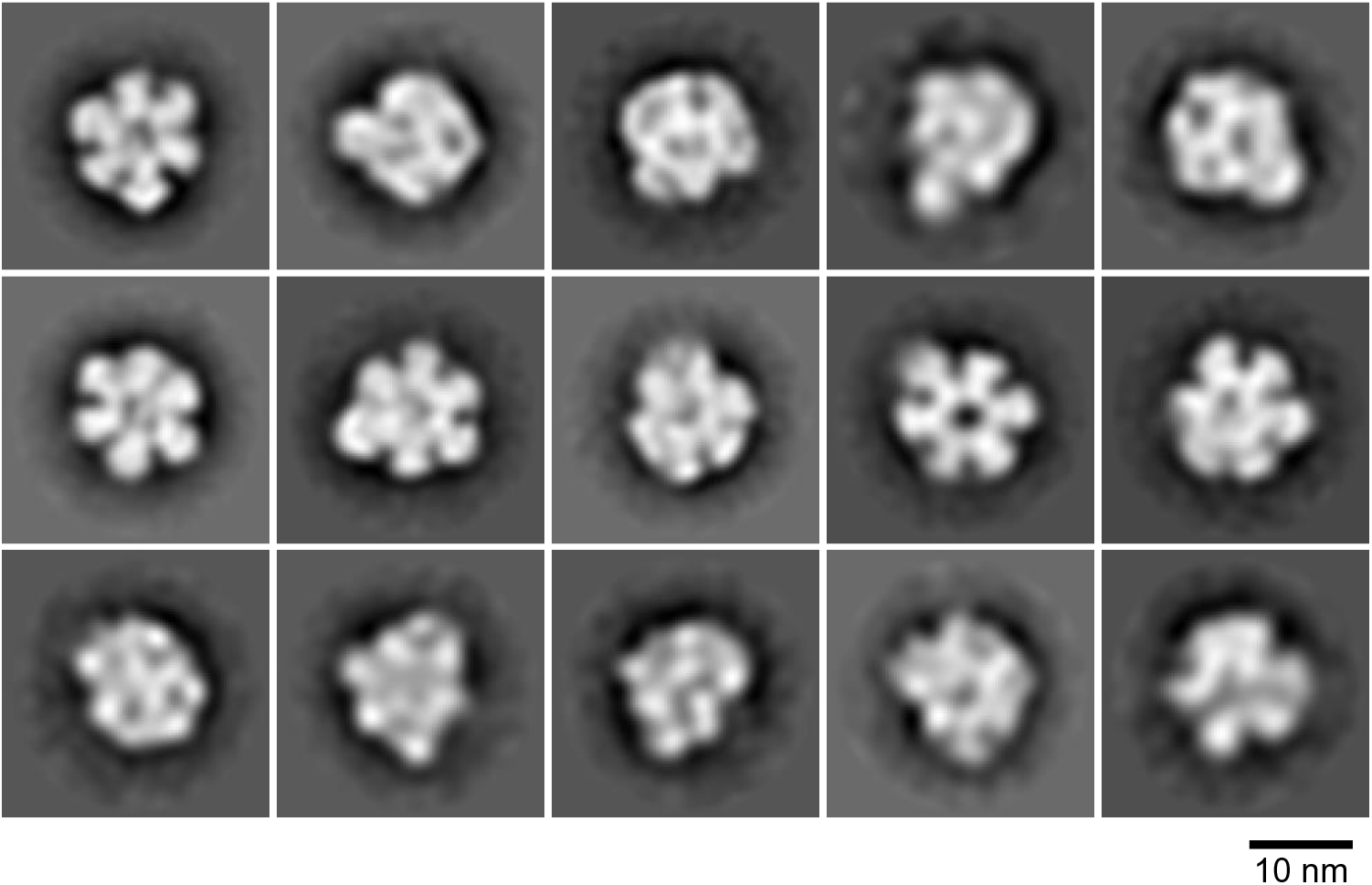
Two-dimensional averaged images of globular complex in Monomer fraction. Fifteen classes of clear particle images from 50 classes are represented.

**FIG S6.**
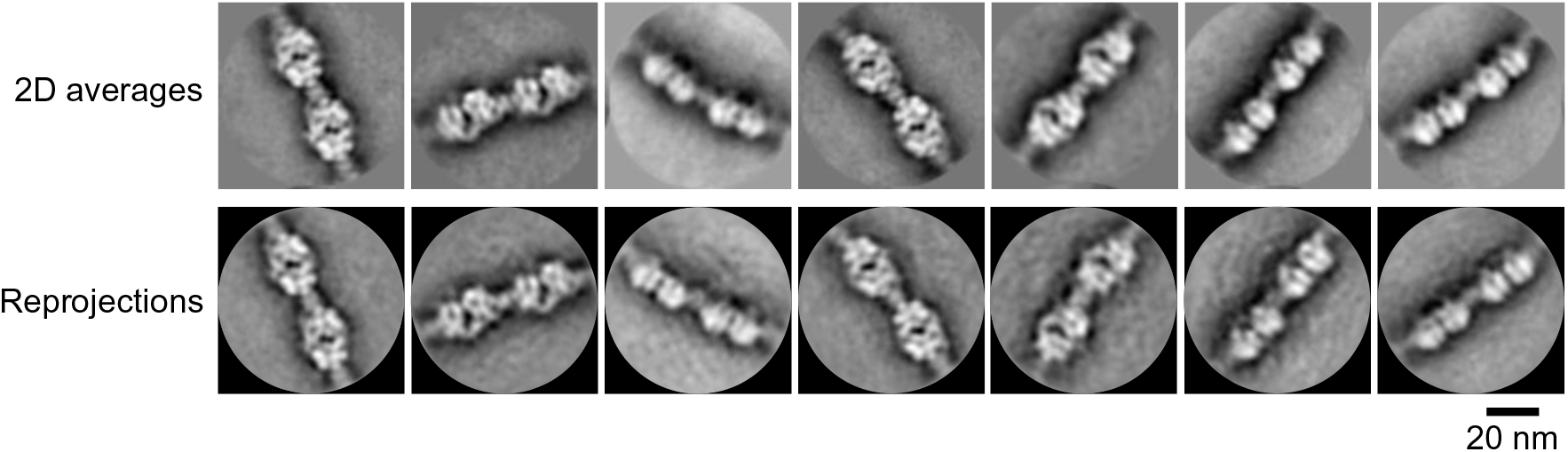
Reprojection images of Chain. Two-dimensional averaged images (upper) and the corresponding reprojection images (lower) calculated from the 3D map of Chain are compared.

**Movie S1 HS-AFM movie showing pattern I particles** The particles were scanned at 10 fps. The scanning field was 70 × 70 nm^2^ with 56 × 56 pixels. The video was played at 10 fps.

**Movie S2 HS-AFM movie showing pattern II particles** The particles were scanned at 10 fps. The scanning field was 70 × 70 nm^2^ with 56 × 56 pixels. The video was played at 10 fps.

**Movie S3 HS-AFM movie showing the shedding process of the peaks.** The particles were scanned at 10 fps. The scanning field was 70 × 70 nm^2^ with 56 × 56 pixels. The video was played at 10 fps. The peaks are indicated by the red triangles.

**Movie S4 HS-AFM movie showing fluctuations in protrusions.** The particles were scanned at 2 fps. The scanning field was 120 × 120 nm^2^ with 120 × 120 pixels. The video was played at 1 fps. The protrusions are indicated by the red triangles.

